# Divergent MIC of Clarithromycin and Azithromycin Against Clinical Isolates of *Mycobacterium abscessus* Complex

**DOI:** 10.1101/2025.05.19.654969

**Authors:** Pusheng Xu, Dan Cao, Yanghui Xiang, Xu Dong, Xiuzhi Jiang, Zhongkang Ji, Yi Li, Xin Yuan, Yuwei Qiu, Kefan Bi, Yiru Zhang, Yuxin Han, Kaijin Xu, Ying Zhang

## Abstract

Current treatment guidelines for *Mycobacterium abscessus* complex (MAB) infections recommend macrolides (clarithromycin/CAM and azithromycin/AZM) as cornerstone therapies, considering them clinically interchangeable without distinction. This study evaluated the comparative in vitro activity of clarithromycin (CAM) and azithromycin (AZM) against 146 clinical isolates of *Mycobacterium abscessus* complex organisms. MIC distributions revealed significantly higher resistance to AZM (94.5% resistant at ≥8 μg/mL) compared to CAM (77.4% resistant at ≥8 μg/mL), with only 4.8% of isolates susceptible to both drugs. Notably, 26 strains (17.8% of total) showed AZM resistance despite being non-resistant to CAM (MIC <8 μg/mL) among 146 clinical isolates. Among the 33 CAM-non-resistant isolates (including susceptible and intermediate strains), 78.8% (26/33) were AZM-resistant, demonstrating frequent discordance in susceptibility profiles. These findings contrast with current treatment guidelines that recommend using CAM and AZM interchangeably without distinction. Subspecies analysis showed three distinct susceptibility profiles: isolates susceptible to both antibiotics (n=3, *M. massiliense*), isolates resistant to both antibiotics (n=3, *M. massiliense*), and *M. bolletii* with CAM-susceptible/AZM-resistant phenotypes (n=2). This differential resistance pattern suggests that while both drugs face substantial resistance challenges, CAM retains meaningful activity against certain strains that are AZM-resistant-particularly in *M. bolletii*. These findings indicate that AZM and CAM should not be used interchangeably to treat MAB infections and emphasize the need to prioritize CAM over AZM in treatment regimens.

## INTRODUCTION

The *Mycobacterium abscessus* complex (MABC), comprising *M. abscessus* subsp. *abscessus, M. massiliense*, and *M. bolletii*, is an emerging group of rapidly growing nontuberculous mycobacteria (NTM) increasingly associated with chronic pulmonary infections, particularly in individuals with underlying lung diseases such as cystic fibrosis and bronchiectasis (1,2). These infections are notoriously difficult to treat due to MABC’s intrinsic resistance to most antibiotics, including β-lactams and aminoglycosides, as well as its ability to rapidly develop acquired resistance during therapy (2,3). Current treatment recommendations focus on combination regimens typically including a macrolide (clarithromycin or azithromycin) as the cornerstone agent, combined with amikacin clofazimine, imipenem and cefoxitin (1). Management of MABC infections poses a major clinical challenge, often requiring prolonged, multidrug treatment regimens with high toxicity and suboptimal success rates (4–6). The overall cure rate for MAB infections remains low, below 50%, due to inherent antibiotic resistance, persistence of the organism, and the absence of clinically effective drugs (7,8).

Macrolides, particularly clarithromycin (CAM) and azithromycin (AZM), have long served as the cornerstone of MABC treatment due to their ability to inhibit protein synthesis by binding to the 50S ribosomal subunit (9). Griffith et al. emphasized that therapy should be guided by in vitro susceptibility testing while noting the essential role of macrolides. However, their efficacy is significantly influenced by the presence of *erm(41)*-mediated inducible resistance, which varies by subspecies: functional *erm(41)* genes in *M. abscessus subsp. abscessus* and *M. bolletii* confer inducible macrolide resistance, while *M. massiliense,* which lacks functional *erm(41)*, has no inducible resistance (10,11). This necessitates either phenotypic (14-day macrolide incubation) or genotypic (line probe assay) confirmation of resistance for non-massiliense isolates (1,10). The 2020 Clinical Practice Guidelines for the Treatment of Nontuberculous Mycobacterial Pulmonary Disease did not differentiate between the efficacy of clarithromycin and azithromycin, considering them interchangeably in the clinical treatment of *Mycobacterium abscessus* (7). The guidelines for diagnosis and trearment of non-tuberculous mycobacteria disease (2020 edition) also did not distinguish between clarithromycin and azithromycin in the use of macrolides in the treatment of *Mycobacterium abscessus*. However, these recommendations were largely based on pharmacologic advantages rather than robust comparative efficacy data, as direct head-to-head clinical trials between these macrolides are lacking. Current treatment regimens typically combine a macrolide (either clarithromycin or azithromycin) with amikacin and one or more additional agents (such as cefoxitin, imipenem, or tigecycline). Importantly, susceptibility to macrolides has been consistently associated with favorable treatment outcomes, highlighting the critical need for accurate susceptibility testing. Howerver, current treatment guidelines do not differentiate between CAM and AZM in terms of drug selection, primarily due to limited comparative susceptibility data. This knowledge gap is particularly acute given that in vitro susceptibility testing forms the basis for treatment decisions in refractory cases (1) and have an impact on treatment outcomes (11). A previous study with *M. avium-intracellulare* indicated that some strains show preferential activity to CAM over AZM, but so far, there have been no studies on comparative testing of AZM and CAM activity in MABC (12).

This study aims to directly compare the in vitro activity of CAM and AZM against a large collection of clinical MABC isolates, analyze their resistance profiles, and investigate potential differences in susceptibility patterns among MABC subspecies. By applying uniform breakpoints for comparative purposes and incorporating molecular subspecies identification, we provide much-needed evidence to inform macrolide selection in clinical practice. Our findings challenge current assumptions about macrolide interchangeability of AZM and CAM and highlight the urgent need for updated treatment guidelines that account for differential resistance patterns between these critically important antibiotics for the treatment of MABC infections.

## RESULTS

### MIC distributions of clarithromycin and azithromycin for MABC

We determined the minimum inhibitory concentrations (MICs) of clarithromycin and azithromycin against 146 clinical isolates of MABC organisms. The susceptibility profiles revealed striking differences between these two macrolides. For clarithromycin, concentrations showed a bimodal distribution with peaks at >128 μg/mL (59 isolates, 40.4%) and 2-32 μg/mL (52 isolates, 35.6%). Azithromycin demonstrated a markedly different profile, with 125 isolates (85.6%) clustering at the highest concentrations (≥128 μg/mL). The remaining 21 isolates (14.4%) showed scattered distribution between 1-64 μg/mL, with no discernible concentration peaks (Fig 1). The AZM MIC values were significantly higher than the CAM MIC values (*P* < 0.001, paired *t*-test) among 146 clinical isolates. These results clearly indicate that clinical isolates of *Mycobacterium abscessus* complex exhibit significantly higher resistance rates to azithromycin compared to clarithromycin. Notably, when compared to the reported peak serum concentration Cmax of clarithromycin (3 μg/mL), 20 clinical isolates (13.7%) exhibited MICs below this threshold, suggesting potential clinical efficacy for these strains. In stark contrast, none of the isolates demonstrated MICs below the Cmax (0.4 μg/mL) (13) of azithromycin, further highlighting clarithromycin’s superior in vitro activity. These findings carry important clinical implications as current treatment guidelines for *M. abscessus* infections do not differentiate between azithromycin and clarithromycin when recommending macrolide-based regimens. Moreover, the alarmingly high prevalence of macrolide-resistant strains (with 76.7% of isolates resistant to both agents) presents a formidable challenge for clinical management.

**Fig 1.**
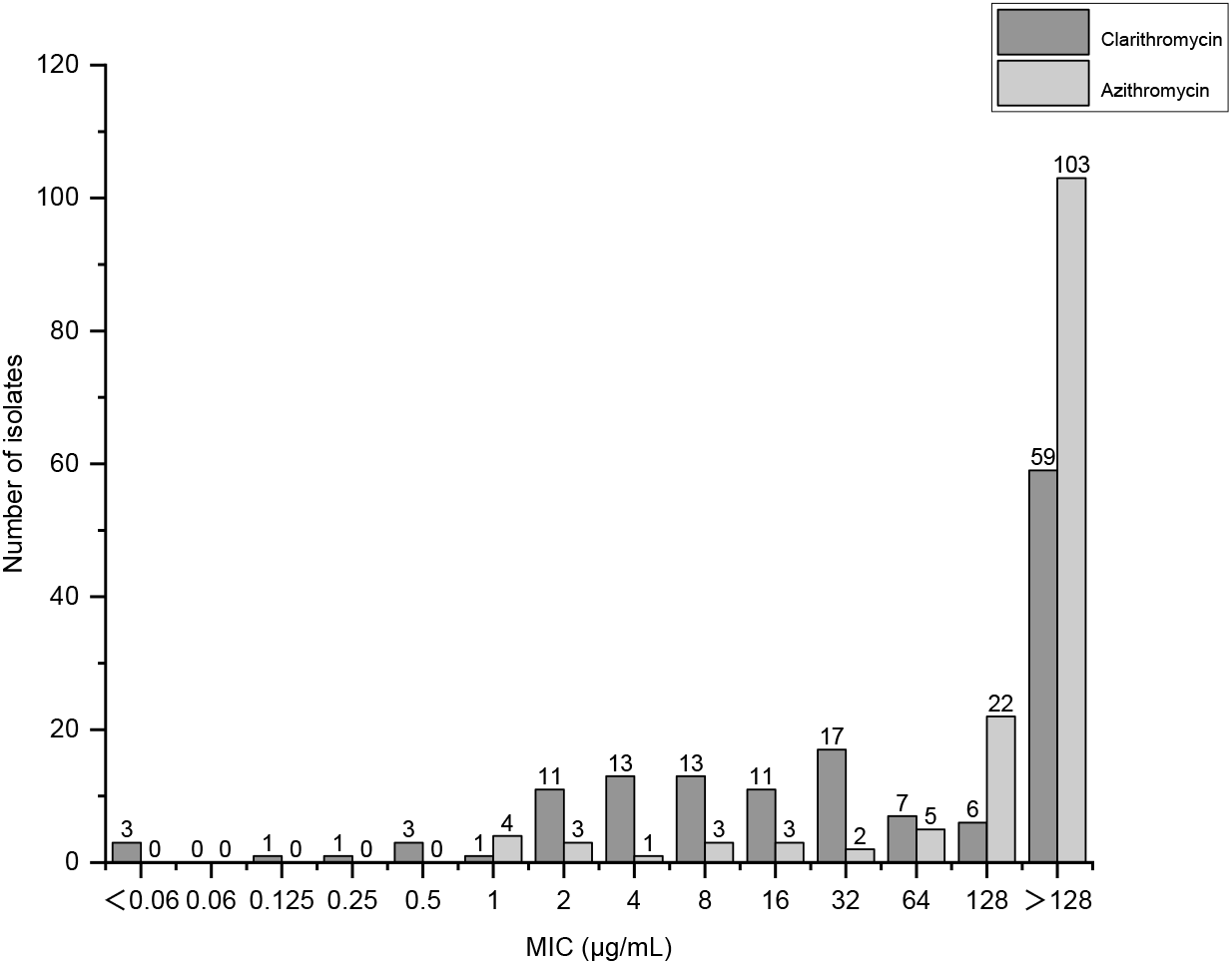
Distribution of clarithromycin and azithromycin MICs. The dark gray bars represent the clarithromycin MICs, and the light gray bars represent the azithromycin MICs.

### Relationship between CAM and AZM susceptibility patterns

The susceptibility profiles of 146 *M. abscessus* clinical isolates to clarithromycin and azithromycin were analyzed based on CLSI interpretive criteria. The established clinical resistance breakpoint for CAM is ≥8 μg/mL and the guidelines’ assumption of therapeutic interchangeability between macrolides (7). Notably, the reference strain ATCC 19977 exhibited a typical susceptibility pattern: sensitive to clarithromycin with MIC =2 μg/mL but resistant to azithromycin with MIC ≥128 μg/mL, which aligned with the overall resistance trends observed in clinical isolates. To facilitate comparative analysis, we uniformly applied a resistance threshold of ≥8 μg/mL to both antibiotics in this study. When examining susceptibility categories more closely, the data showed that 113 isolates (77.4%) met resistance criteria for CAM (≥8 μg/mL), with 20 strains (13.7%) classified as susceptible (MIC ≤2 μg/mL) and 13 (8.9%) as intermediate (4 μg/mL). In contrast, AZM resistance was even more prevalent, affecting 138 isolates (94.5%), with only 7 susceptible (4.8%) and 1 intermediate (0.7%) (Table 1). Analysis of resistance patterns revealed significant differences between the two macrolides. Among all tested isolates, 112 strains (76.7%) demonstrated resistance to both antibiotics, while only 7 isolates (4.8%) remained susceptible to both agents. Subspecies analysis of these 7 dual-susceptible isolates showed a distinct distribution: 3 belonged to *M. massiliense* and 4 to *M. abscessus subsp. abscessus*. Notably, among 33 CAM susceptible/intermediate isolates, 26 strains were resistant to AZM, accounting for 78.7% (26/33) of the isolates. By contrast, the reverse pattern (CAM-resistant/AZM-susceptible) was exceptionally rare, occurring in just one isolate (0.7%) (Fig 2). Detailed analysis of these 26 AZM-resistant isolates revealed extreme resistance profiles: 18 strains (69.2%) exhibited MICs ≥128 μg/mL, while the remaining showed graded resistance - 2 strains (7.7%) at 64 μg/mL, 1 strain (3.8%) at 32 μg/mL, 3 strains (11.5%) at 16 μg/mL, and 2 strains (7.7%) at 8 μg/mL. This distribution underscores that even among CAM-susceptible M. abscessus isolates, high-level AZM resistance predominates, with nearly 70% reaching the maximum measurable MIC concentration. Figure 3 shows the relationship between the AZM MIC and CAM MIC for each isolate. The Pearson correlation coefficient of AZM-MIC and CAM-MIC was 0.3264 (95% CI: 0.1731–0.4642, *P* < 0.001).

**Table 1.** Distribution of Susceptibility, Intermediate, and Resistance Categories for Clarithromycin and Azithromycin Against *Mycobacterium abscessus* Clinical Isolates.

| Antibiotics | Susceptible | Intermediate | Resistant |
| --- | --- | --- | --- |
| Clarithromycin | 20 | 13 | 113 |
| Azithromycin | 7 | 1 | 138 |

**Fig 2.**
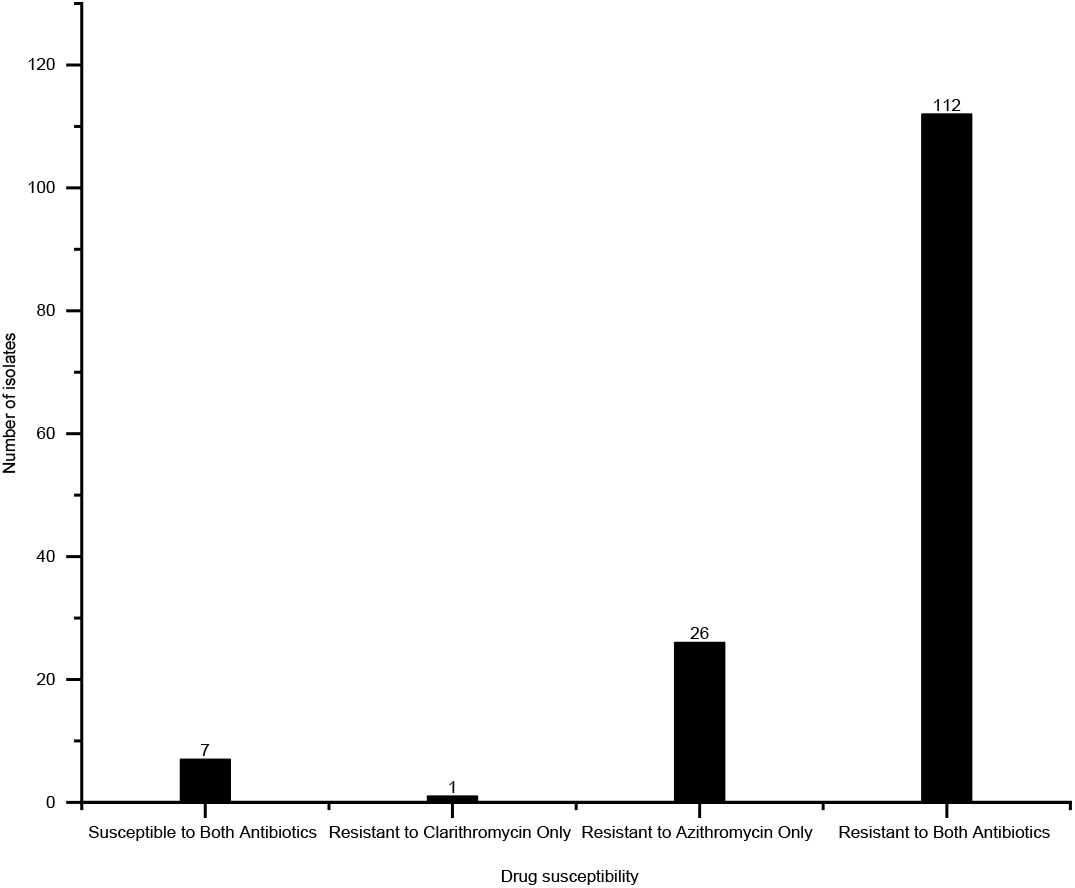
Comparison of the sensitivity of azithromycin and clarithromycin. Different columns reflect different sensitivity profiles of clinical strains to azithromycin and clarithromycin.

**Fig 3.**
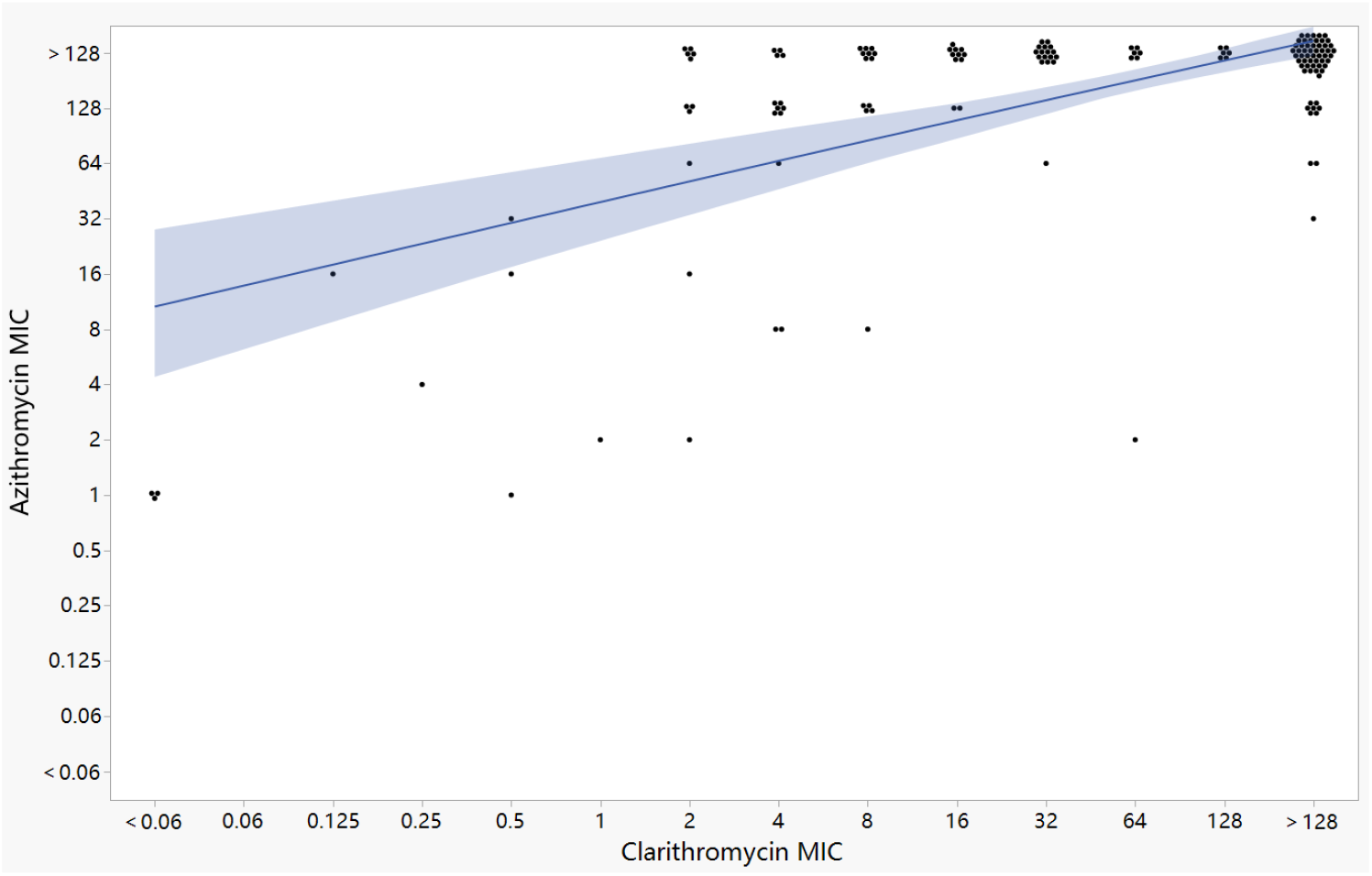
Correlation between the clarithromycin and azithromycin MICs. Scatter dot plots showing the relationship between the clarithromycin and azithromycin MICs. Each dot represents one isolate. The regression line and 95% CI are shown in blue.

Cross-resistance was common (76.7% resistant to both) which indicates partial overlap of resistance mechanisms. Despite no official AZM breakpoint, these findings support prioritizing CAM in treatment, though susceptibility testing remains critical. Standardized AZM clinical breakpoints are needed for accurate resistance assessment.

### Subspecies-specific differences in MIC profiles

The analysis of MIC distributions revealed significant subspecies-dependent variations in susceptibility patterns among the MABC isolates. Three distinct phenotypic profiles emerged: *M. abscessus subsp. massiliense* isolates exhibited either complete macrolide susceptibility (MIC ≤1 μg/mL for both drugs, n=3) or pan-resistance (MIC >128 μg/mL, n=3); Both *M. abscessus subsp. bolletii* isolates demonstrated the unusual clarithromycin-susceptible/azithromycin-resistant profile (CAM MIC=0.5 μg/mL; AZM MIC=16-32 μg/mL). This tripartite resistance distribution suggests multiple concurrent resistance mechanisms at play. The complete susceptibility of sensitive *M. massiliense* isolates aligns with its characteristic *erm(41)* gene truncation (9), while their resistant counterparts likely acquired additional resistance mutations. The discordant CAM-susceptible/AZM-resistant profile in *M. bolletii* implies either differential efflux pump activity or ribosomal target site variations specific to azithromycin. Particularly noteworthy is the correlation between subspecies identity and susceptibility pattern, emphasizing the necessity of subspecies-level identification for clinical decision-making.

## DISCUSSION

Our study reveals a critical divergence in the in vitro activity of clarithromycin (CAM) and azithromycin (AZM) against MABC clinical isolates, with AZM demonstrating significantly higher resistance rates (94.5%) compared to CAM (77.4%). Notably, 17.8% of isolates exhibited AZM resistance despite CAM susceptibility, while the reverse pattern was rare (0.7%). Of the 33 *M. abscessus* clinical isolates exhibiting clarithromycin susceptibility/intermediate, 26 (78.8%) were resistant to azithromycin (MIC ≥8 μg/mL). These findings challenge the current guideline assumption of interchangeable AZM and CAM use and suggest that CAM retains superior activity over AZM against certain MABC strains, particularly *M. bolletii*. Subspecies-level analysis further highlighted distinct resistance profiles, with *M. massiliense* isolates displaying either pan-susceptibility or pan-resistance, while *M. bolletii* isolates uniquely exhibited CAM-susceptible/AZM-resistant phenotypes. These results underscore the necessity of individualized susceptibility testing and subspecies identification to guide macrolide selection in clinical practice.

The mechanisms underlying the observed differential resistance patterns warrant further investigation. The higher AZM resistance may stem from its weaker ribosomal binding affinity or greater susceptibility to efflux pumps (14), potentially due to fundamental structural differences between the two macrolides (Fig 4). AZM’s 15-membered lactone ring structure, compared to CAM’s 14-membered ring, along with its additional amino sugar moiety, may contribute to reduced target binding and enhanced recognition by efflux systems like MmpL4/5 in MABC. These structural variations could also influence membrane permeability and intracellular drug accumulation (Fig 4). Future research should employ whole-genome sequencing and transcriptomic analyses to elucidate these molecular determinants, particularly focusing on mutations in 23S rRNA domain V and efflux pump regulation. Furthermore, standardized clinical breakpoints for AZM against MABC are urgently needed to harmonize resistance interpretation and improve treatment guidelines, especially considering these pharmacodynamic differences. The structural basis of differential resistance could be further investigated through advanced techniques like cryo-EM to visualize drug-ribosome interactions at atomic resolution.

**Fig 4.**
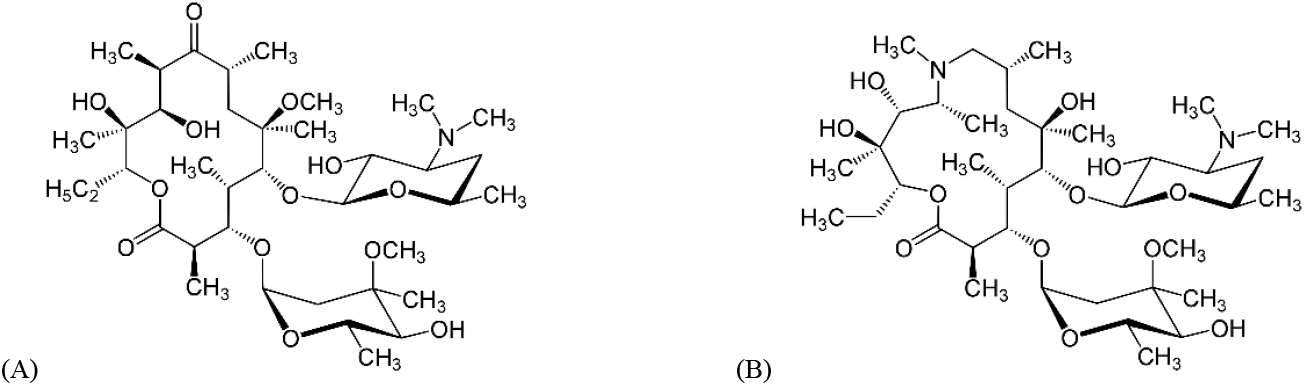
Structures of Azithromycin (A) and Clarithromycin (B). Azithromycin and clarithromycin differ structurally in their lactone ring size (15-membered vs 14-membered) and functional groups. Azithromycin lacks the C-3 cladinose sugar and contains an extra amino sugar, while clarithromycin retains cladinose with a C-6 methoxy group.

Clinically, our findings advocate for a paradigm shift in MABC management. While AZM’s pharmacokinetic advantages have made it a preferred choice (15), its inferior in vitro activity suggests that CAM should be more reliable in empiric regimens, especially for *M. bolletii* infections. However, the high prevalence of pan-resistant strains (76.7%) underscores the need for novel therapeutic strategies, such as macrolide potentiators (e.g., efflux pump inhibitors) (16) or repurposed non-macrolide agents (e.g., bedaquiline or clofazimine). Prospective clinical trials comparing CAM- and AZM-based regimens, stratified by subspecies and resistance profiles, are essential to validate our in vitro observations and refine treatment algorithms in future studies.

In conclusion, this study highlights the inadequacy of treating CAM and AZM as interchangeable in therapies for MABC infections. The stark resistance disparities, coupled with subspecies-specific patterns, emphasize the importance of routine susceptibility testing and molecular identification in guiding macrolide selection. While CAM appears to be the more reliable option while considering MIC values, the pervasive resistance to both agents calls for expanded research into alternative therapies and resistance mechanisms. Our findings suggest the need for future guideline revisions to optimize treatment outcomes in this challenging patient population.

## MATERIALS AND METHODS

### Bacterial growth and culture conditions

This retrospective cross-sectional study collected all strains at The First Affiliated Hospital, Zhejiang University School of Medicine (Hangzhou, China). All strains were characterized and identified as *M. abscessus complex* through 16S rRNA gene sequencing analysis. Strains were grown in Middlebrook 7H9 medium (Becton Dickinson) supplemented with 10% oleic acid-dextrose-catalase (OADC, Aladdin) and 0.05% vol/vol Tween 80 (Sigma-Aldrich).

### Determination of MICs

CAMHB broth (Hopebio) was used to prepare twofold dilutions of Clarithromycin, Azithromycin (Macklin), ranging from 128 μg/ml to 0.0625 μg/ml. Both compounds were dissolved in DMSO. The stationary phase cultures were diluted to about 10^6^ CFU/ml, added to antibiotic containing 96-well plate, which was wrapped with tin foil and placed in a 37°C incubator and visible bacterial growth was observed after 5 days.

### Statistical analysis

A paired *t*-test was performed to compare the logarithms of AZM and CAM MIC distribution and the Pearson correlation coefficient was calculated to assess their correlation. All statistical analyses were performed using JMP and Origin.

## ACKNOWLEDGMENTS

This study was supported by a National Infectious Disease Medical Center startup fund (YZ)(B2022011-1), and Jinan Microecological Biomedicine Shandong Laboratory project (JNL-2022050B).

